# Structure and position-specific interactions of prion-like domains in transcription factor Efg1 phase separation

**DOI:** 10.1101/2023.11.09.566450

**Authors:** Szu-Huan Wang, Tongyin Zheng, Nicolas L. Fawzi

## Abstract

*Candida albicans*, a prominent member of the human microbiome, can make an opportunistic switch from commensal coexistence to pathogenicity accompanied by an epigenetic shift between the white and opaque cell states. This transcriptional switch is under precise regulation by a set of transcription factors (TFs), with Enhanced Filamentous Growth Protein 1 (Efg1) playing a central role. Previous research has emphasized the importance of Egf1’s prion-like domain (PrLD) and the protein’s ability to undergo phase separation for the white-to-opaque transition of *C. albicans*. However, the underlying molecular mechanisms of Efg1 phase separation have remained underexplored. In this study, we delved into the biophysical basis of Efg1 phase separation, revealing the significant contribution of both N-terminal (N) and C-terminal (C) PrLDs. Through NMR structural analysis, we found that Efg1 N-PrLD and C-PrLD are mostly disordered though have prominent partial α-helical secondary structures in both domains. NMR titration experiments suggest that the partially helical structures in N-PrLD act as hubs for self-interaction as well as Efg1 interaction with RNA. Using condensed-phase NMR spectroscopy, we uncovered diverse amino acid interactions underlying Efg1 phase separation. Particularly, we highlight the indispensable role of tyrosine residues within the transient α-helical structures of PrLDs particularly in the N-PrLD compared to the C-PrLD in stabilizing phase separation. Our study provides evidence that the transient α-helical structure is present in the phase separated state and highlights the particular importance of aromatic residues within these structures for phase separation. Together, these results enhance the understanding of *C. albicans* TF interactions that lead to virulence and provide a crucial foundation for potential antifungal therapies targeting the transcriptional switch.

**Statement of Significance:** Phase separated condensates have been found across the domains of life and many types of cells. To understand their varied functions, seeing the residue-by-residue details of the structure and interactions of component protein constituents is essential. A set of transcription factors that phase-separate controls cell fate of the pathogenic yeast Candida albicans. Here, we examine the structural and interaction details of a main regulator of this process, Efg1, using NMR spectroscopy and biochemical assays. We find Efg1’s phase-separating domains are not entirely disordered as often assumed but in fact contain helical regions that persist upon phase separation. We also reveal the balance of contacts formed in the condensed phase and the importance of specific residues and regions in phase separation.

## Introduction

Known as one of the most common commensal fungi in the human microbiome, *Candida albicans* is also an opportunistic pathogen that can cause potentially life-threatening infections and pose a significant threat to human health (1, 2). The virulence of *C. albicans* is intricately linked to its ability to undergo phenotypical switching between white and opaque cell states (2, 3). This epigenetic transition is under the tight regulation of a transcriptional network, wherein the protein Enhanced Filamentous Growth Protein 1 (Efg1) plays a pivotal role (1, 4–7). Additionally, Efg1 modulates crucial processes like hyphae formation and filamentation (8–11). Moreover, it stands at the core of biofilm formation (11, 12), a vital aspect of *C. albicans* pathogenicity, making Efg1 a promising target for anti-fungal therapies.

Recent studies have unveiled the intriguing phenomenon of phase separation among transcription factors (TFs), especially Efg1, leading to the formation of biomolecular condensates within cells (6, 13). These condensates play a critical role in governing fungal cell states. The prion-like domains (PrLDs) of Efg1 are indispensable for its function in the white-opaque switch and likely mediate interactions between TFs (6). However, the precise mechanisms through which these PrLDs mediate Efg1 self-interaction and phase separation remain unprobed. Although a few studies have examined low complexity domain phase separation using residue-by-residue details from NMR spectroscopy, the protein domains examined thus far have primarily been human proteins with distinct sequence composition from Efg1 (14–18). Additionally, the domains studied so far have been essentially disordered without significant population of partial secondary structure and hence do not represent the entire range of transient structures formed by intrinsically disordered domains that phase separate. Yet, previous studies have established the pivotal role of transient α-helical secondary structure in mediating protein self-interaction and phase separation in the case of the human RNA-binding protein TDP-43 (19, 20) or where dispersed phase oligomerization of helices opposes phase separation in the case of the glutamine-rich RNA-binding proteins from the multinucleate fungus *Ashbya gossypii* (21). Hence, it is important both to identify secondary structure in phase separating proteins and evaluate secondary structure contribution to phase separation. Furthermore, no studies have directly visualized secondary structure with residue-by-residue resolution in condensed phases to help understand how these structures play a role in phase separation.

In this study, we delve into the biophysical basis of Efg1 phase separation, focusing on its PrLDs. We demonstrate that both PrLDs, Efg1 N and Efg1 C, exhibit substantial disorder with transient secondary structure features. Using solution NMR techniques, we illustrate the diversity of intermolecular contacts between amino acid types that drive the phase separation process. Importantly, our investigation reveals a crucial role for tyrosine residues within the transient α-helical structures of PrLDs. By deciphering the underlying biophysical principles governing Efg1 phase separation, this research sheds light on the intricate molecular mechanisms regulating *C. albicans* virulence, potentially paving the way for targeted therapeutic interventions against fungal infections.

## Material and Methods

### Protein expression and purification

The full-length Efg1 protein, fused with a histidine-tagged maltose binding protein (MBP) and TEV protease cleavage site (his-MBP-FL Efg1), was cloned into the pTHMT vector and subsequently transformed into BL21 cells. For non-labeled protein expression, the transformed cells were cultured in Luria-Bertani (LB) medium, whereas for isotopically labeled protein, M9 minimal medium supplemented with appropriate isotopes was utilized. Cultures were grown at 37°C with continuous shaking until reaching an optical density at 600 nm (OD600) of 0.6-0.8. Expression of the fusion protein was induced by adding 1 mM Isopropyl β-D-1-thiogalactopyranoside (IPTG) and incubating the cultures at 37°C for 4 hours. Following induction, the bacterial cells were harvested and lysed in the lysis buffer (20mM Tris pH8, 500mM NaCl pH8) using a sonicator to disrupt the cell membranes. The lysed cell suspension was centrifuged, and the resulting supernatant containing the His-MBP-FL Efg1 protein was collected. To purify the protein from cell lysate, a 5 ml HisTrap HP column was employed. The sample-loaded HisTrap column was washed with 20 mM Tris pH 8, 1 M NaCl, 10mM imidazole buffer, then the bound protein was eluted with 20 mM Tris pH 8, 1 M NaCl, 300 mM Imidazole buffer. Subsequently, the protein was further purified using a Superdex 200 (26 600) size-exclusion column, yielding a highly purified protein sample ready for further analysis.

Efg1 N-PrLDB, N-PrLDB, or NC-PrLD, fused with a histidine-tag followed by TEV protease cleavage site, were cloned into the pJ411 vector and transformed into BL21 cells. The expression protocol is similar to that of FL-MBP-Efg1 described above. The harvested cells were resuspended and lysed in the lysis buffer (20mM Tris pH8, 500mM NaCl pH8) using a sonicator. The insoluble fraction containing the expressed proteins was collected and solubilized in a denaturing buffer composed of 20 mM Tris pH 8, 8 M Urea, 500 mM NaCl, and 10 mM imidazole. This urea-containing buffer effectively solubilized the inclusion bodies. The solubilized proteins were purified using a Histrap column. The column-bound proteins were washed with 20 mM Tris pH 8, 8 M Urea, 500 mM NaCl, and 10 mM imidazole buffer to remove non-specifically bound impurities. The purified proteins were eluted from the column using the elution buffer containing 20 mM Tris pH 8, 8 M Urea, 500 mM NaCl, and 300 mM imidazole. The purified protein was then buffer exchanged into 20 mM CAPS buffer pH 11 for storage (tyrosine deprotonates at pH 11, leading to disruption of phase separation and aggregation) (16).

### Phase separation quantification

The phase separation behavior of tag-free Efg1-PrLDs was assessed by diluting the protein from a 20 mM CAPS, pH 11.0, stock solution to a concentration of 200 μM into 50 mM Tris, pH 7.4, supplemented with the appropriate salt concentration. Samples were then centrifuged at 14,000 g for 10 minutes at room temperature. All experiments were conducted in triplicate. The protein concentration in the remaining supernatant was quantified using a NanoDrop spectrophotometer. Extinction coefficients were predicted from the primary sequence by ProtParam (Expasy).

To quantify FL-Efg1 phase separation, MBP-tagged protein was treated with TEV protease, targeting the TEV cleavage site positioned between MBP and the Efg1 protein. The mixture was incubated at 30°C for 3 hours to ensure complete cleavage. Following the incubation period, the sample was centrifuged at 14,000g for 10 minutes to separate the phases. The absorbance at 280 nm (A280) of the supernatant was measured. To obtain accurate results, the absorbance corresponding to TEV protease was subtracted from the measured A280. The decrease in A280 absorbance after subtraction indicated the amount of FL-Efg1 that had undergone phase separation after liberation from the N-terminal solubilizing MBP fusion, and the remaining amount of soluble Efg1 can be calculated by subtracting phase-separated Efg1 from the total protein amount.

### NMR sample preparation and NMR spectroscopy

Isotopically labeled protein samples of Efg1 PrLD were prepared in a buffer containing 80 mM MES pH 6.2, 100 mM NaCl, and 5% D_2_O. The condensed phase Efg1 N-PrLD samples were generated by diluting stock protein in 80 mM MES 100mM NaCl 10% D2O buffer, torula yeast RNA was also added to stabilize the condensed phase at 4:1 protein-to-RNA ratio (by mass). The phase-separated protein solution was then allowed to settle at 4 °C overnight to form a macroscopic condensed phase. The macroscopic condensed phase was then moved into a 3 mm NMR tube using a glass pipette for the collection of NMR experiments.

NMR spectra were recorded on Bruker Avance 600MHz or 850 MHz ^1^H Larmor frequency spectrometers with HCN TCl z-gradient cryoprobes. Two-dimensional ^1^H-^15^N HSQCs were acquired using spectral widths of 12.0 ppm and 22.0 ppm in the direct and indirect dimensions, with 4096 and 512 total points and acquisition time of 240 ms and 135 ms, respectively. ^1^H-^13^C HSQC spectra were acquired using spectral widths of 12.0 ppm and 76.0 ppm in the direct and indirect dimensions, with 4096 and 400 total number of points and acquisition time of 200 ms and 12 ms, respectively.

Triple resonance experiments, including HNCACB, CBCA(CO)NH, HNCO, HNCA, and HNN, were conducted to establish sequential and side-chain resonance assignments. Acquisition times for the triple resonance experiments were 185 ms in the direct ^1^H dimensions, 24 ms in the indirect ^15^N dimensions, 15 ms in the indirect C’ dimensions and 6 ms in the indirect C_α_/ C_β_ dimensions. Spectral widths were 13 ppm in ^1^H, 22 ppm in ^15^N, 7.5 ppm in C’ dimension and 54 ppm in C_α_/C_β_ dimension. Data processing and analyses followed standard protocols using NMRPipe and CCPNMR Analysis 2.5 software (22, 23).

Motions of the backbone of Efg1 N-PrLD and C-PrLD were probed by ^15^N spin relaxation experiments at 850 MHz ^1^H Larmor frequency (20 Tesla) using standard pulse sequences (hsqct1etf3gpsitc3d, hsqct2etf3gpsitc3d, hsqcnoef3gpsi). ^15^N *R*_2_ experiments were performed with 256 and 2,048 total points with acquisition times of 74 ms and 115 ms and a sweep width of 10.5 and 20 ppm centered at 117 and 4.7 and ppm in the indirect ^15^N and direct ^1^H dimensions, respectively. *R*_2_ experiments consist of six interleaved relaxation delays with an interscan delay of 2.5 s, and total *R*_2_ relaxation CPMG loop lengths of 16.5, 264.4, 33.1, 132.2, 66.1, 198.3 ms. ^15^N *R*_1_ experiments were recorded with 256 and 3072 total points with acquisition times of 74 ms and 115 ms and a sweep width of 20 and 10.5 ppm centered at 117 and 4.7 ppm in the indirect 15N and direct ^1^H dimensions, respectively. Each ^15^N R1 experiment was made up of six interleaved ^15^N *R*_1_ relaxation delays of 5, 1,000, 100, 800, 500, and 300 ms. (^1^H) ^15^N heteronuclear NOE (hetNOE) experiments were made up of interleaved sequences with and without proton saturation, with a recycle delay of 5 s. (^1^H) ^15^N hetNOE experiments were recorded with 256 and 4,096 total points with a sweep width of 20 and 10.5 ppm centered at 117 and 4.7 ppm in the indirect ^15^N and direct ^1^H dimensions, respectively.

Intermolecular NOE-based experiments were recorded on condensed Efg1 N-PrLD containing a 50:50 mixture of ^15^N/^13^C (∼99% isotopic incorporation) and ^14^N/^12^C Efg1 N-PrLD (natural abundance), or entirely ^15^N/^13^C Efg1 N-PrLD as a control. Three-dimensional ^13^C/^15^N filtered NOESY-^13^C (edited)-HSQC experiments were recorded with a mixing time of 250 ms and with 64, 128, and 4,096 total points with sweep widths of 80, 12, and 12 ppm centered at 43, 4.7, and 4.7 ppm for aliphatic regions in the F3 dimension or sweep widths of 45, 12, and 12 ppm centered at 122.5, 4.7, and 4.7 ppm for aromatic regions in the F3 dimension, for indirect ^13^C, indirect ^1^H, and direct ^1^H dimensions, respectively. We note that 250 ms mixing time showed no signs of significant spin diffusion in similar phases of FUS disordered domains (24).

## Results

### Efg1 prion-like domains drive phase separation

Previous studies identified the N-terminal and C-terminal regions of Efg1 as having prion-like sequence character and are essential for Efg1 phase separation (6). Secondary structure predictions based on the amino acid primary sequence predict that Efg1 contains two intrinsically disordered, prion-like domains in both the C- and N-terminus and a structured DNA binding domain (DBD) (Fig. 1A). Despite the long evolutionary distance suggesting there is no homology (i.e. common ancestor), the amino acid composition of the Efg1 PrLDs resembles that of low complexity (LC) disordered domains from other phase-separation capable proteins such as Fused in Sarcoma that readily phase separates (16, 25) (Fig. 1B). However, Efg1 PrLDs have a lower fraction of S and Y residue content, a higher fraction of Q and N content, and do have some aliphatic residues including I, L, and V as well as F aromatic residues that are all completely absent in FUS LC. Hence the sequences are similar but even compositionally show differences.

**Figure 1.**
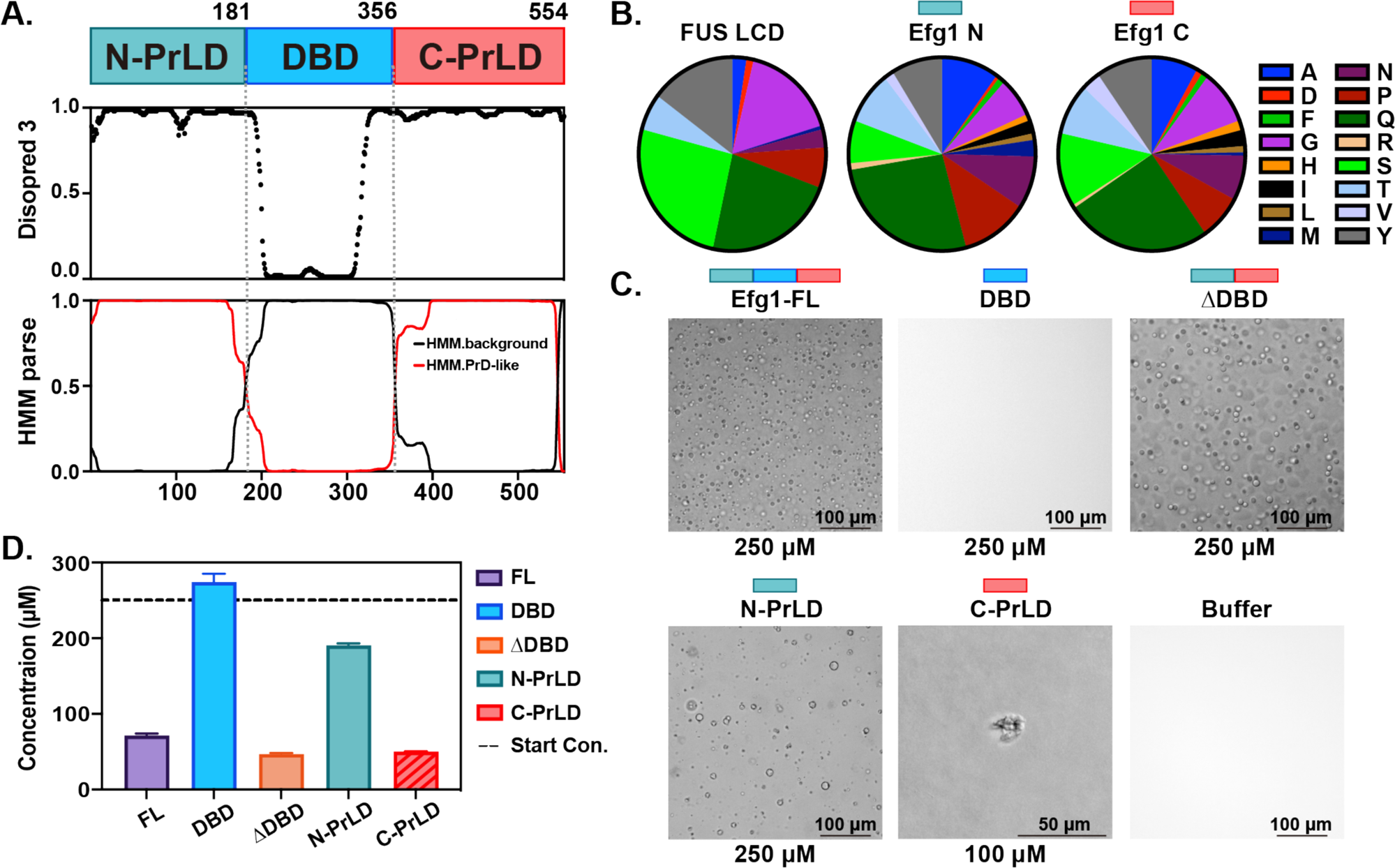
Efg1 prion-like domains drive phase separation. A) Efg1 contains two disordered prion-like domains (PrLD), one on the N-terminal end and one on the C-terminal end, flanking a central DNA binding domain (DBD). Both the N- and C-PrLDs are predicted to be disordered and prion-like, while the structured DBD is a conserved KilA-N DNA binding domain. B) Comparison between the amino acid compositions of FUS low complexity domain, Efg1 N-PrLD and Efg1 C-PrLD. The three sequences have similar compositions though the fraction of serine and tyrosine are lower in Efg1 while the fraction of aliphatic residues is higher in Efg1. C) Differential interference contrast microscopy shows that the full-length Efg1, as well as ΔDBD and N-PrLD, can undergo phase separation to form liquid-like droplets. The isolated DBD remains soluble at tested conditions while the C-PrLD forms aggregates. D) The saturation concentration for phase separation of Efg1 full-length protein measured by the concentration of Efg1 protein remaining in the supernatant after phase separation followed by centrifugation to sediment droplets and/or aggregates. Full-length and ΔDBD have similar saturation concentrations (*C*_sat_), while the N-PrLD alone phase separates less. The C-PrLD shows a low apparent *C*_sat_ but this value is complicated by the formation of solid assemblies, indicated by the diagonal striped fill.

To elucidate the contribution of each Efg1 domain to phase separation, we conducted a series of experiments involving various Efg1 domain deletion and isolation constructs. First, we examined the phase separation behavior of the full-length (FL) Efg1 and compared that to deletion of the DNA-binding domain, Efg1 ΔDBD, and the isolated DBD. The isolated DBD did not exhibit phase separation on its own, nor does its presence augment the phase separation of full-length Efg1 compared to the Efg1 ΔDBD (i.e. N-PrLD fused to C-PrLD) constructs (Fig. 1C,D). These results strongly suggest that the DBD does not play a direct role in facilitating phase separation.

Next, we investigated the phase separation propensity of the individual PrLDs of Efg1 (N-PrLD and C-PrLD). Unlike the DBD, isolated N-PrLD demonstrated liquid-like phase separation behavior, forming droplets in solution, albeit at a higher protein saturation concentration (*c*_sat_) compared to the FL protein. In contrast, the C-terminal region of Efg1 (Efg1 C-PrLD) did not form liquid droplets, instead forming gel-like aggregates. Nevertheless, the observed solubility difference between Efg1 N and NC-PrLD suggests that the C-terminal region still contributes to Efg1 phase separation (Fig. 1D), potentially influencing the process through interactions with the N-terminal domain. Hence, though the N- and C-PrLDs have similar sequence composition, and are both prone to self-assemble and both contribute to the observed phase separation of the full-length Efg1, they have distinct material properties at these solution conditions.

### Efg1 prion-like domains contain transient α-helical structures

To identify potential secondary structures in the PrLDs of Efg1, we analyzed Efg1 primary sequence using the program PSIPRED (26), which predicted the presence of α-helical structure in both PrLDs (Fig. 2B). To validate the presence of transient helical structures within the Efg1 PrLDs, we obtained NMR chemical shift assignments of backbone N, H_N_, C’ (i.e. carbonyl CO), C_α_, and sidechain C_β_ positions for both domains using triple resonance NMR approaches (see Methods). The presence of partially populated helical structure in specific regions of both Efg1 N-PrLD and Efg1 C-PrLD is evident based on sensitive NMR-based measures including positive values of the difference in residual C_α_ and C_β_ chemical shift positions compared to a random coil reference, ΔδC_α_ – ΔδC_β_, and the SSP (secondary structure propensity score) calculated from these chemical shifts (Fig. 2C,D) (27), as well as high helix scores in δ2D analysis (Fig. S1) (28). As transient α-helical structure will also slow local reorientational motions that can be detected by backbone ^15^N NMR spin relaxation experiments (19), we found signatures of slower motions *R*_1_, *R*_2_, and heteronuclear NOE that corroborate the existence of slower-moving secondary structures in the sequences (Fig. 2E and Fig. S2). Among the helical regions in N-PrLD and C-PrLD, we found that two polyQ stretches within Efg1 N and one within the Efg1 C domain adopted partial α-helical conformations (Fig. 2C).

**Figure 2.**
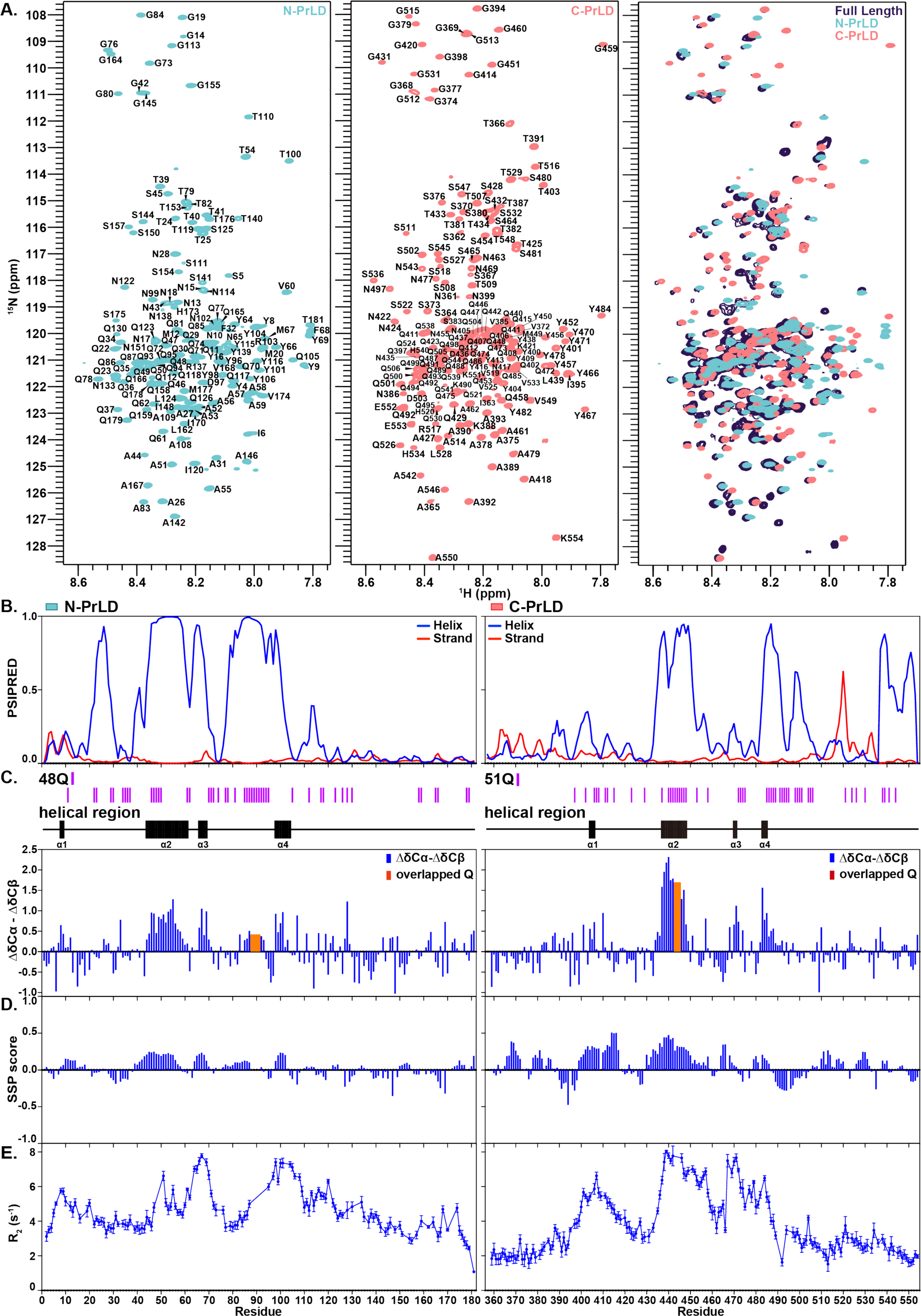
The two prion-like domains in Efg1 are largely disordered, with transient helical structure features. A) ^1^H-^15^N HSQC spectra of isolated N-PrLD, C-PrLD, and the two overlayed on FL-Efg1. Backbone resonances from the two PrLDs are mostly present in the spectra of full-length Efg1. Resonances in the spectra of the isolated domain overlap well with the full-length protein spectrum suggesting no major structural changes resulting from protein truncation. B) Secondary structure prediction based on Efg1 Primary sequence using the program PSIPRED. C) NMR experimental secondary chemical shifts and D) NMR chemical shift-based secondary structure analysis using the program SSP indicates the formation of transient α-helices in both the N-PrLD and C-PrLD.

To assess whether truncation affected the overall structure of the individual domains, we compared the NMR spectra of the isolated N and C domains to the corresponding regions within the FL Efg1 protein. As we expected, the NMR peaks from both isolated domains remained largely unchanged from those observed in the FL protein spectrum, indicating that truncation did not significantly alter the overall structural integrity of the two domains (Fig. 2A).

### Helical regions in Efg1 N-PrLD are hubs for RNA interaction and self-interaction

Given the significance of the Efg1 prion-like domains (PrLDs) in driving self-interaction and phase separation, we conducted a comprehensive examination of prion-like domain self-interactions using NMR. We focused our investigations on Efg1 N-PrLD as C-PrLD did not form stable liquid forms under the tested conditions, and both PrLDs displayed similar sequence compositions and secondary structure characteristics.

To identify residues in Efg1 that mediate self-interaction, we performed concentration titration experiments at concentrations below the *c*_sat_ threshold for phase separation. Through these titrations, we observed chemical shift perturbations (CSPs) (Fig. 3A), as well as decreased peak intensity (Fig. 3B) at the helical regions within the Efg1 N-PrLD. As observed for the helical region embedded in the prion-like TDP-43 C-terminal domain (19, 20, 29), these findings indicate that these transient helical conformations within Efg1 N-PrLD may contribute to self-interaction in the dispersed phase (i.e. before phase separation).

**Figure 3.**
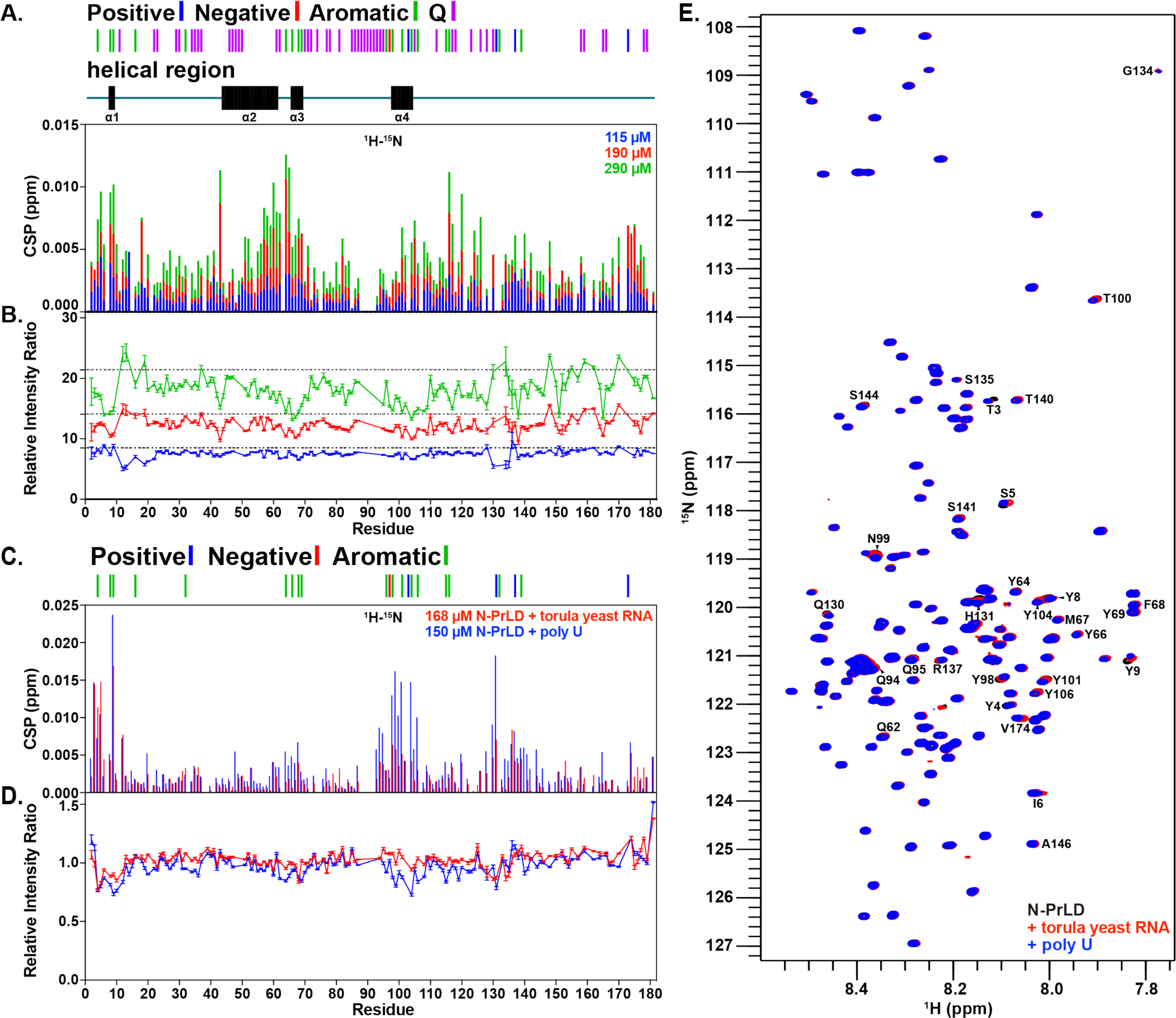
Helical regions in Efg1 N-PrLD mediate transient N-PrLD self-interaction and RNA contacts. A-B) Consistent with transient interactions, the four helical regions in Efg1 N-PrLD see A) increased chemical shift perturbations (CSP) and B) decreased relative peak intensity ratios compared to the rest of the protein, as the result of increased protein concentration. Data are relative to a low concentration reference spectrum at 13.5 μM. C-D) The helical regions also exhibit C) chemical shift and D) relative peak intensity ratios perturbation induced by the addition of either torula yeast or poly U RNA, suggesting that the regions also mediate non-specific RNA interaction. E) ^1^H-^15^N HSQC spectra overlay of N-PrLD and N-PrLD with the addition of either torula yeast or poly U RNA. The most prominent peak shifts are observed at the N-terminus and at residues in the fourth helical region.

Considering that Efg1 functions as a transcription factor that may use disordered regions to interact with RNA (30), we further hypothesized that its PrLDs may also engage in non-specific interactions with RNA. To test this, we carried out NMR titration experiments using Efg1 N-PrLD and two types of RNAs, an unstructured RNA (poly U) and an RNA extract with highly heterogenous structures (torula yeast RNA). The NMR CSPs and peak intensities after addition of these RNAs are remarkably similar, suggesting that Efg1 N-PrLD interacts with RNA in a non-sequence-specific manner (Fig. 3C,D). Strikingly, the sites of RNA interaction coincided with the helical regions within the protein sequence, which are also implicated in self-interaction. This dual interaction profile of Efg1 N-PrLD with RNA and self-interaction is intriguing and suggests a possible interplay between RNA binding and phase separation.

### Helical Regions Mediate Interaction in Efg1 N-PrLD Condensed Phase

To gain further insights into the molecular interactions driving Efg1 phase separation, we created macroscopic condensed phase samples by first inducing phase separation in solution and then centrifuging the droplets to form a separate phase (Fig. 4B). Interestingly, we found that these macroscopic condensates “aged”, changing from liquid-like to gel-like over a matter of hours. Given that we found RNA interactions with Efg1, we reasoned that RNA may help solubilize Efg1. By including a modest amount of RNA (∼25% of the mass of the protein) in the phase separation reaction, we found that the samples remained liquid-like for weeks. Using these samples of Efg1 N-PrLD with RNA for high-resolution NMR, we compared the condensed phase spectra with those obtained in the dispersed phase. We observed overall spectral overlap in ^1^H-^15^N and especially ^1^H-^13^C HSQC spectra (Fig. 4A, Fig. S3) of condensed phase and dispersed phase Efg1 N-PrLD, supporting the notion that the overall primarily disordered structure of the protein remains largely similar in both phases, as seen for the disordered domains of FUS (16, 18).

**Figure 4.**
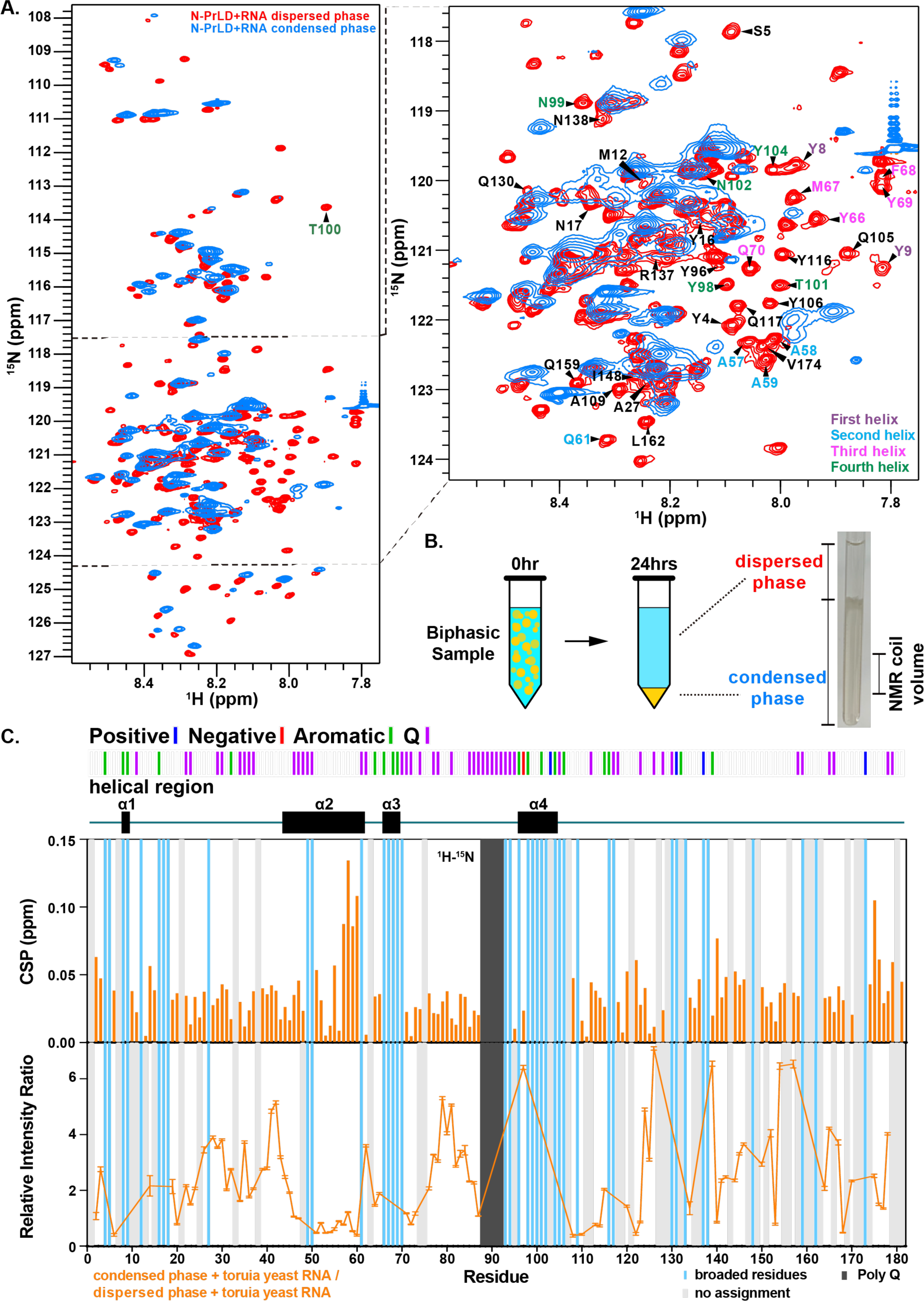
The helical regions mediate contacts in Efg1 N-PrLD condensed phase. A) ^1^H-^15^N HSQC spectra overlay of the dispersed phase and condensed phase Efg1 N-PrLD. The majority of residues remain in similar positions in the two phases, suggesting a similar overall conformation. B) The Efg1 condensed NMR samples were made by first inducing phase separation in a conical tube, then allowing the droplets to settle and form a condensed phase over 24 hours. The condensed phase can then be transferred into an NMR tube for measurements. C) Chemical shift perturbations and relative intensity ratios between Efg1 N-PrLD ^1^H-^15^N HSQC spectra of the condensed phase and the dispersed phase suggest that the second helical region in N-PrLD participates in molecular interaction that stabilizes the condensed phase. Peak broadening (blue) observed in the first, third, and fourth helical regions indicate their contribution to mediating interactions in the condensed phase.

Previous studies on macroscopic condensed phases of disordered regions from several groups have uniformly demonstrated that the disordered regions show little change in the spectra upon phase separation (14–16, 24). However, these studies focused on proteins that lacked any significant population of secondary structure. Intriguingly, we find that residues within the helical regions of Efg1 N-PrLD exhibited substantial peak broadening beyond detection in the condensed phase, while the rest of the protein displayed similar relative NMR signals in both phases (Fig. 4C). This observation indicates that the helical regions either experience slower motion in the condensed phase compared to the rest of the protein or undergo chemical exchange caused by self-interactions during Efg1 phase separation.

Moreover, we mapped out chemical shift perturbations (CSPs) and intensity ratios of the two phases (Fig. 4C), and identified significant CSPs in the second helical region, along with peak broadening in the first, third, and fourth helices. Alanine residues in the second helix A57/A58A/59) form a cluster of resonances in the dispersed phase (Fig. 2A, left) that may correspond to the resonances shifted more up field (i.e. to the right and up) than other resonances in the spectra of the condensed phase (Fig. 4A, right), although the fast spin relaxation of these resonances in the condensed phase have precluded confirmation of the assignment. In the dispersed phase, these alanine residue positions give rise to C_β_ resonances characteristic of α-helical positions (Fig. S3). Importantly, these alanine C_β_ resonances with α-helical chemical shifts remain present in the condensed phase, suggesting that α-helical structure is retained after phase separation of Efg1 N-PrLD. Together, these finding suggest that helical regions within Efg1 N-PrLD are present in condensed phases and participate in phase-separation-driving self-interaction, possibly as structural elements acting as key “interaction hubs” in Efg1.

### Intermolecular Contacts Mediated by Amino Acid Types in Efg1 Phase Separation

To probe the nature of intermolecular interactions during Efg1 phase separation, we created isotopically mixed-labeled macroscopic condensed phases (mixing equal amounts of fully ^13^C/^15^N protein with unlabeled protein) and employed filter/edited NOESY experiments (Fig. 5A) that therefore report only on intermolecular contacts that are formed in the condensed phase (18, 31). To ensure the accuracy of our analyses, we prepared a separate sample entirely made of fully labeled protein, enabling us to subtract NOE intensities stemming from imperfect isotopic incorporation of NMR active isotopes as well as incomplete NOE filtering along with other artifacts. Our investigations encompassed sidechain-sidechain and sidechain-backbone interactions (Fig. 5B,C, and Fig. S4), offering insights into the specific amino acid types involved in mediating intermolecular contacts during phase separation. Although it is not possible to interpret these NOE intensities quantitatively on an absolute scale, it is instructive to compare the intensity of the various pairs. Notably, tyrosine and glutamine residues exhibited the highest NOE intensities (Fig. 5C), suggesting a significant contribution of these residues to the interaction network, similar to previously reported data on FUS (24). Additionally, hydrophobic residues such as alanine, isoleucine, leucine, valine, and proline also exhibit significant NOE intensities suggesting a high contribution of hydrophobic interactions in Efg1 phase separation. Interestingly, intermolecular NOEs between glutamine (H_γ_) and alanine are more prominent than tyrosine (H_γ_) and alanine, suggesting some residue pair contact preferences. These slight preferences are also apparent when the NOE intensities are normalized by the residue frequency in Efg1 N-PrLD (Fig. S4B).

**Figure 5.**
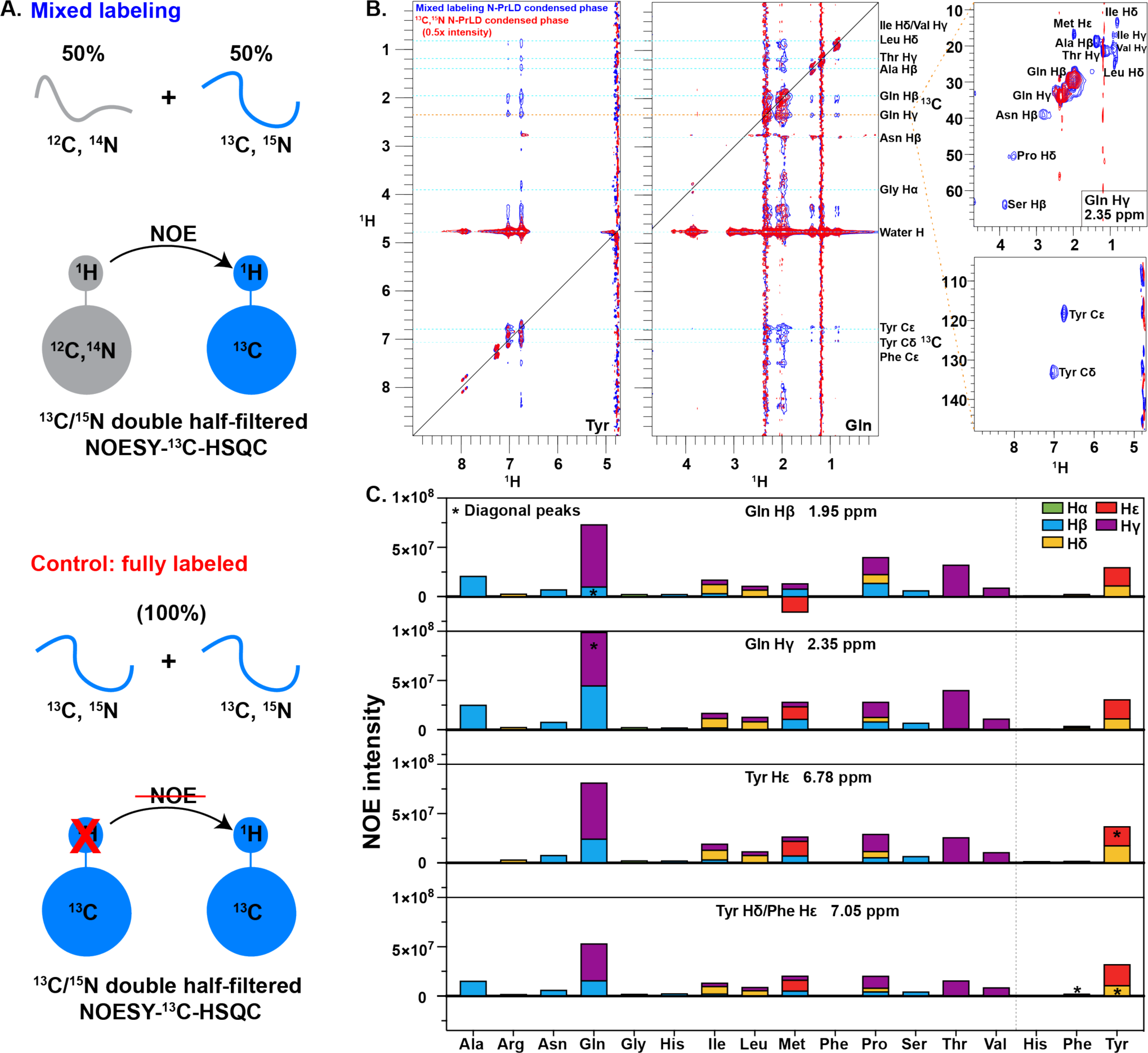
Intermolecular contacts between many residue types in Efg1 phase separation. A) Equimolar amounts of ^13^C/^15^N double-labeled and non-labeled Efg1 N-PrLD were used to create a condensed phase NMR sample in order to conduct filter/edited HSQC experiments to probe intermolecular interactions in the phase. The control experiment was carried out using a condensed phase sample consisting of only ^13^C/^15^N double-labeled N-PrLD. B) ^1^H-^1^H 2D projection of the ^13^C/^15^N double-half filtered NOESY-^13^C HSQC (left) and the NOE stripe for Gln H_γ_ (right). Artifacts are removed by subtracting ½ peak height of the fully labeled control (red) from the mixed labeled experiment (blue). C) Quantification of intermolecular NOE from non-labeled Gln H_β_, Gln H_γ_, Tyr H_ε_, and Tyr H_δ_/Phe H_ε_ to ^13^C-attached protons on protein sidechains. The NOE intensities for ^1^H resonances attached to aliphatic and aromatic ^13^C carbon were collected in separate experiments, as indicated by the dashed line separating the two sets of values.

### Tyrosine Residues in Helical Regions Stabilize Phase Separation

With our analysis of the structures and contacts formed by Efg1 PrLDs in phase separation, we then sought to investigate their precise role in phase separation using mutagenic approaches. First, we engineered proline mutations within the helical segments of Efg1 N to break the structures and assess impacts (Fig. 6A). Through NMR secondary chemical shift analysis and ^1^H ^15^N HSQC chemical shift perturbations, we confirmed the successful disruption of the helical structures (Fig. S5). These mutations produced quantitative reduction in the phase behavior of Efg1 (Fig. 6A). Though the change in the apparent saturation concentration for phase separation was small, the effects of multiple simultaneous mutations were mostly additive, consistent with a model where interactions are formed by multiple helical regions in each Efg1 PrLD. We note that Efg1 ΔDBD phase separation is enhanced by increased sodium chloride concentration, suggesting a role of hydrophobic interactions in driving phase separation. In comparison to FUS LCD, which also demonstrates ‘salt out’ phase behavior (24), Efg1 ΔDBD exhibits lower sensitivity to sodium chloride. This disparity could be attributed to Efg1’s elevated charged amino acid composition and the involvement of electrostatic interactions. The presence of additional salt weakens these electrostatic interactions, counteracting the augmented hydrophobic interaction, thereby impact of the added salt on phase separation within the range of physiological salt concentrations is reduced.

**Figure 6.**
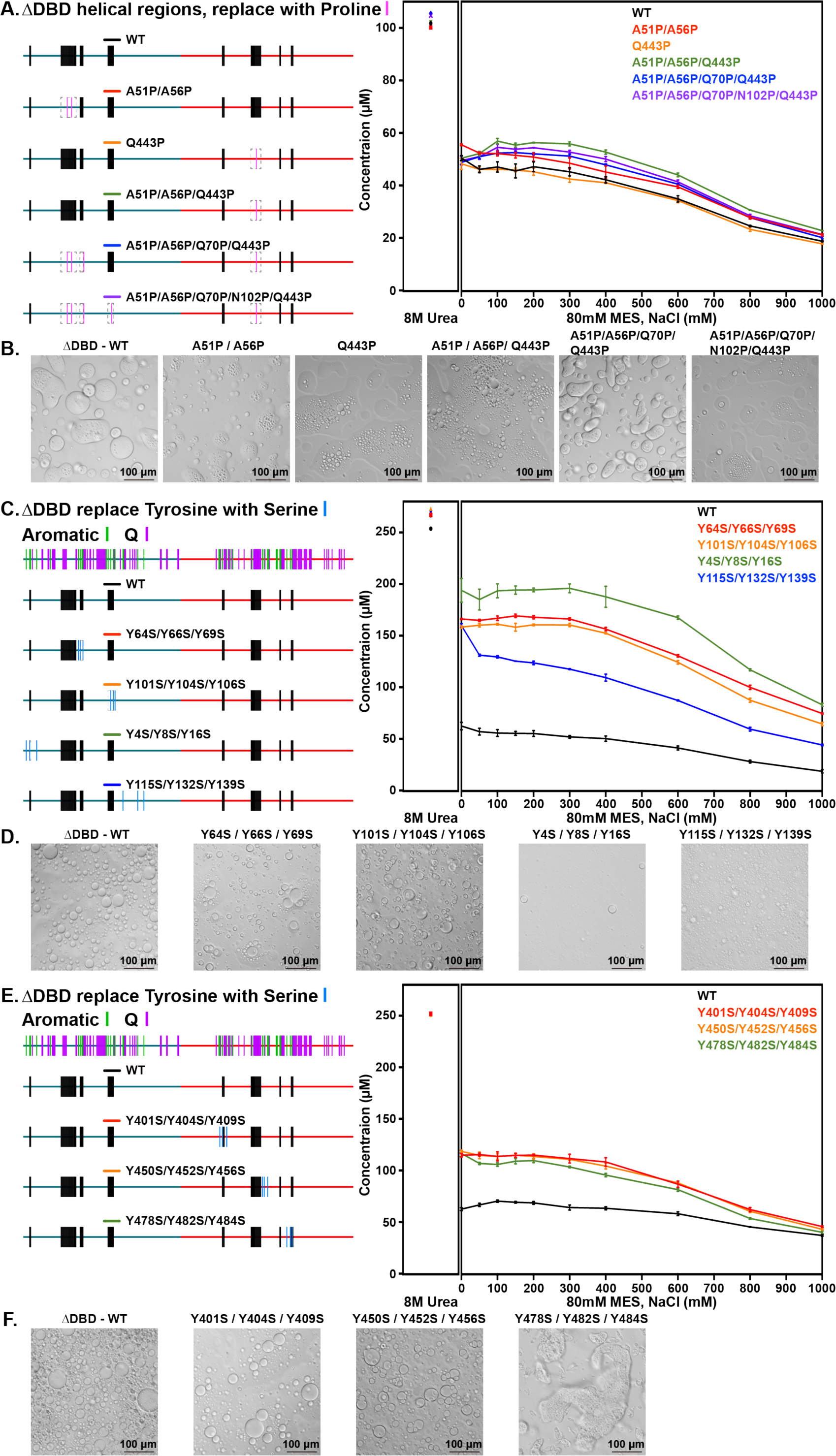
Tyrosine residues in the helical regions of Efg1 N-PrLD are especially important for phase separation. A-B) Proline mutations that disrupt helical structures in Efg1 ΔDBD do not significantly alter the phase behavior of the protein. C-D) Tyrosine mutations in the N-PrLD helices more drastically reduce the phase separation propensity of Efg1 ΔDBD compared to mutations in the disordered regions, as shown by the significantly increased saturation concentrations of the helical tyrosine mutations. E-F) Tyrosine mutations in the C-PrLD helices and disordered regions both reduce phase separation of Efg1 ΔDBD to similar extents. Mutations in the last C-PrLD helical region cause the protein to form gel-type aggregates.

Drawing from previous studies (32, 33) and our data that both highlighted the critical role of tyrosine residues in promoting phase separation, we then tested the role of tyrosine residues within both the helical and disordered regions of Efg1 N. We introduced mutations that replaced aromatic tyrosine with serine in these regions (34), and subsequently assessed their effects on phase separation (Fig. 6C,E). Remarkably, all mutants exhibited lower phase separation propensities, underscoring the importance of tyrosine residues in mediating Efg1 phase separation. Among the variants, tyrosine to serine substitutions in the helical regions of N-PrLD displayed the most potent effects on phase separation (Fig. 6C), suggesting that tyrosine residues within these N-PrLD structural elements may play a more prominent role in promoting Efg1 phase separation than those outside these secondary structures. Intriguingly, Y-to-S substitutions in the C-PrLD displayed distinct effects compared to their counterparts in N-PrLD in two major ways. Firstly, the substitutions in C-PrLD were generally less potent compared to those in N-PrLD. More importantly, mutations in the C-PrLD did not exhibit the same secondary-structure-dependence seen in N-PrLD. Except for the mutation in the last helical region (Y115S/Y132S/Y139S), which resulted in the formation of gel-like aggregates, mutations in both the helical and disordered regions of C-PrLD showed a similar decrease in phase separation. Taken together, our observation highlights the distinct roles that N-PrLD and C-PrLD play in stabilizing Efg1 phase separation, and provides crucial insights into the specific contributions of N-PrLD helical regions to the stabilization of Efg1 phase separation.

## Discussion

In summary, our study has probed some of the molecular intricacies of the phase separation of Efg1 from *Candida albicans*. We have shown that Efg1 contains two intrinsically disordered prion-like domains, Efg1 N and Efg1 C, with significant disorder and partial secondary structure features. These domains play a crucial role in mediating Efg1 phase separation, where the N-terminal domain is sufficient for droplet formation, the C-terminal domain enhances phase separation into a liquid form even if it forms solid structures on its own. Notably, we identified partially α-helical structures within these domains that remain partially helical upon phase separation. These helical regions have emerged as hubs for RNA interaction and self-interaction, indicating a multifaceted role for Efg1 in cellular processes. The self-interaction mediated by helical structures in Efg1 PrLD aligns with similar observations in proteins such as TDP-43 CTD (19, 20), although we do not find evidence for helical enhancement upon self-assembly as observed for TDP-43 (19). Though the effect on phase separation is small in magnitude, disrupting the α-helices decreases phase separation, unlike the α-helices in Whi3 that drive clustering but discourage phase separation (21). The dynamic and transient nature of the helical regions implied by their partial population might facilitate versatile interactions with both helical structures and disordered regions. Consequently, disrupting these helical structures did not lead to complete phase separation disassembly but instead caused a substantial reduction.

Additionally, our findings emphasize the vital contribution of tyrosine residues in stabilizing Efg1 phase separation, as seen for other proteins including FUS (24, 32, 33). By disrupting these tyrosine-mediated interactions, we attenuated Efg1 phase separation, suggesting a central role for these residues in driving the process. Tyrosine residues in the N-PrLD helical structures had particularly high contribution to phase separation, prompting us to consider the intriguing possibility that the transient helical structures may position the tyrosine side chains in certain orientations that facilitate favorable interactions, driving Efg1 phase separation. These results also suggest that tyrosine residues in each prion-like domain can have a distinct quantitative contribution to phase separation. However, our findings also show that many residue types make contact in Efg1 PrLD phase separation, as observed for FUS (18). Efg1 has a distinct amino acid composition with aliphatic hydrophobic amino acids. Recent data showed that hydrophobic amino acids contribute to PrLD phase separation in TDP-43 (29), but could not asses what contact pairs these residues form. Here we show that hydrophobic Efg1 residue make extensive contacts in the condensed phase, particularly with glutamine residues and to a lesser extent with tyrosine. This suggests that glutamine may directly interact with hydrophobic amino acids, an ability that has been overlooked. This critical relationship between the secondary structure elements and specific amino acid residues adds a new dimension to our understanding of Efg1 phase separation and the underlying molecular mechanisms governing this biophysical phenomenon. These biophysical insights not only enhance our understanding of *Candida*’s pathogenicity but also could pave the way for innovative antifungal therapies that alter phase separation to alter the epigenetic switch that determines cell fate.

## Author Contributions

TZ and NLF designed research. SHW performed research. SHW and TZ analyzed data and created visualizations. TZ and NLF wrote the manuscript.

## Acknowledgements

The authors thank the members of the Fawzi laboratory, Richard Bennett, and Corey Frazer for helpful suggestions. We thank Mandar Naik and the Structural Biology Core Facility at Brown University for assistance and support with NMR infrastructure. Research was supported in part by NSF MCB 1845734 (to N.L.F.), NIGMS R01GM147677 (to N.L.F.), and a Pape Adams Postdoctoral Award from the Carney Institute for Brain Science at Brown University (to TZ).

## Data and resource availability

NMR chemical shift assignments for Efg1 N-PrLD and C-PrLD are deposited at the Biological Magnetic Resonance Database (BMRB, http://www.bmrb.wisc.edu/) (*Deposition identifiers in process*). Plasmids generated for this project are deposited at Addgene.org at https://www.addgene.org/Nicolas_Fawzi/ (*Deposition identifiers in process*). Detailed protocols associated with this study can be obtained by contacting the corresponding authors.

## Reference

1. Glazier, V.E. 2022. EFG1, Everyone’s Favorite Gene in Candida albicans: A Comprehensive Literature Review. Front. Cell. Infect. Microbiol. 12.

2. Sohn, K., C. Urban, H. Brunner, and S. Rupp. 2003. *EFG1* is a major regulator of cell wall dynamics in *Candida albicans* as revealed by DNA microarrays. Mol. Microbiol. 47:89–102.

3. Sasse, C., M. Hasenberg, M. Weyler, M. Gunzer, and J. Morschhäuser. 2013. White-Opaque Switching of Candida albicans Allows Immune Evasion in an Environment-Dependent Fashion. Eukaryot. Cell. 12:50–58.

4. Srikantha, T., L.K. Tsai, K. Daniels, and D.R. Soll. 2000. EFG1 Null Mutants of Candida albicansSwitch but Cannot Express the Complete Phenotype of White-Phase Budding Cells. J. Bacteriol. 182:1580–1591.

5. Zordan, R.E., D.J. Galgoczy, and A.D. Johnson. 2006. Epigenetic properties of white–opaque switching in Candida albicans are based on a self-sustaining transcriptional feedback loop. Proc. Natl. Acad. Sci. 103:12807–12812.

6. Frazer, C., M.I. Staples, Y. Kim, M. Hirakawa, M.A. Dowell, N.V. Johnson, A.D. Hernday, V.H. Ryan, N.L. Fawzi, I.J. Finkelstein, and R.J. Bennett. 2020. Epigenetic cell fate in Candida albicans is controlled by transcription factor condensates acting at super-enhancer-like elements. Nat. Microbiol. 5:1374–1389.

7. Sonneborn, A., B. Tebarth, and J.F. Ernst. 1999. Control of White-Opaque Phenotypic Switching in Candida albicans by the Efg1p Morphogenetic Regulator. Infect. Immun. 67:4655–4660.

8. Sudbery, P.E. 2011. Growth of Candida albicans hyphae. Nat. Rev. Microbiol. 9:737–748.

9. Lu, Y., C. Su, X. Mao, P.P. Raniga, H. Liu, and J. Chen. 2008. Efg1-mediated Recruitment of NuA4 to Promoters Is Required for Hypha-specific Swi/Snf Binding and Activation in Candida albicans. Mol. Biol. Cell. 19:4260–4272.

10. Braun, B.R., and A.D. Johnson. 2000. TUP1, CPH1 and EFG1 Make Independent Contributions to Filamentation in Candida albicans. Genetics. 155:57–67.

11. Ramage, G., K. VandeWalle, J.L. López-Ribot, and B.L. Wickes. 2002. The filamentation pathway controlled by the Efg1 regulator protein is required for normal biofilm formation and development in Candida albicans. FEMS Microbiol. Lett. 214:95–100.

12. McCall, A.D., R.U. Pathirana, A. Prabhakar, P.J. Cullen, and M. Edgerton. 2019. Candida albicans biofilm development is governed by cooperative attachment and adhesion maintenance proteins. Npj Biofilms Microbiomes. 5:1–12.

13. Staples, M.I., C. Frazer, N.L. Fawzi, and R.J. Bennett. 2023. Phase separation in fungi. Nat. Microbiol. 8:375–386.

14. Brady, J.P., P.J. Farber, A. Sekhar, Y.-H. Lin, R. Huang, A. Bah, T.J. Nott, H.S. Chan, A.J. Baldwin, J.D. Forman-Kay, and L.E. Kay. 2017. Structural and hydrodynamic properties of an intrinsically disordered region of a germ cell-specific protein on phase separation. Proc. Natl. Acad. Sci. 114:E8194–E8203.

15. Reichheld, S.E., L.D. Muiznieks, F.W. Keeley, and S. Sharpe. 2017. Direct observation of structure and dynamics during phase separation of an elastomeric protein. Proc. Natl. Acad. Sci. 114:E4408–E4415.

16. Burke, K.A., A.M. Janke, C.L. Rhine, and N.L. Fawzi. 2015. Residue-by-Residue View of In Vitro FUS Granules that Bind the C-Terminal Domain of RNA Polymerase II. Mol. Cell. 60:231–241.

17. Ryan, V.H., G.L. Dignon, G.H. Zerze, C.V. Chabata, R. Silva, A.E. Conicella, J. Amaya, K.A. Burke, J. Mittal, and N.L. Fawzi. 2018. Mechanistic view of hnRNPA2 low complexity domain structure, interactions, and phase separation altered by disease mutation and arginine methylation. Mol. Cell. 69:465–479.e7.

18. Murthy, A.C., W.S. Tang, N. Jovic, A.M. Janke, D.H. Seo, T.M. Perdikari, J. Mittal, and N.L. Fawzi. 2021. Molecular interactions contributing to FUS SYGQ LC-RGG phase separation and co-partitioning with RNA polymerase II heptads. Nat. Struct. Mol. Biol. 28:923–935.

19. Conicella, A.E., G.H. Zerze, J. Mittal, and N.L. Fawzi. 2016. ALS Mutations Disrupt Phase Separation Mediated by α-Helical Structure in the TDP-43 Low-Complexity C-Terminal Domain. Structure. 24:1537–1549.

20. Conicella, A.E., G.L. Dignon, G.H. Zerze, H.B. Schmidt, A.M. D’Ordine, Y.C. Kim, R. Rohatgi, Y.M. Ayala, J. Mittal, and N.L. Fawzi. 2020. TDP-43 α-helical structure tunes liquid–liquid phase separation and function. Proc. Natl. Acad. Sci. 117:5883–5894.

21. Seim, I., A.E. Posey, W.T. Snead, B.M. Stormo, D. Klotsa, R.V. Pappu, and A.S. Gladfelter. 2022. Dilute phase oligomerization can oppose phase separation and modulate material properties of a ribonucleoprotein condensate. Proc. Natl. Acad. Sci. 119:e2120799119.

22. Delaglio, F., S. Grzesiek, G.W. Vuister, G. Zhu, J. Pfeifer, and A. Bax. 1995. NMRPipe: a multidimensional spectral processing system based on UNIX pipes. J. Biomol. NMR. 6:277–293.

23. Skinner, S.P., R.H. Fogh, W. Boucher, T.J. Ragan, L.G. Mureddu, and G.W. Vuister. 2016. CcpNmr AnalysisAssign: a flexible platform for integrated NMR analysis. J. Biomol. NMR. 66:111–124.

24. Murthy, A.C., G.L. Dignon, Y. Kan, G.H. Zerze, S.H. Parekh, J. Mittal, and N.L. Fawzi. 2019. Molecular interactions underlying liquid−liquid phase separation of the FUS low-complexity domain. Nat. Struct. Mol. Biol. 26:637–648.

25. Patel, A., H.O. Lee, L. Jawerth, S. Maharana, M. Jahnel, M.Y. Hein, S. Stoynov, J. Mahamid, S. Saha, T.M. Franzmann, A. Pozniakovski, I. Poser, N. Maghelli, L.A. Royer, M. Weigert, E.W. Myers, S. Grill, D. Drechsel, A.A. Hyman, and S. Alberti. 2015. A Liquid-to-Solid Phase Transition of the ALS Protein FUS Accelerated by Disease Mutation. Cell. 162:1066–1077.

26. McGuffin, L.J., K. Bryson, and D.T. Jones. 2000. The PSIPRED protein structure prediction server. Bioinformatics. 16:404–405.

27. Marsh, J.A., V.K. Singh, Z. Jia, and J.D. Forman-Kay. 2006. Sensitivity of secondary structure propensities to sequence differences between α- and γ-synuclein: Implications for fibrillation. Protein Sci. Publ. Protein Soc. 15:2795–2804.

28. Sormanni, P., C. Camilloni, P. Fariselli, and M. Vendruscolo. 2015. The s2D Method: Simultaneous Sequence-Based Prediction of the Statistical Populations of Ordered and Disordered Regions in Proteins. J. Mol. Biol. 427:982–996.

29. Mohanty, P., J. Shenoy, A. Rizuan, J.F. Mercado-Ortiz, N.L. Fawzi, and J. Mittal. 2023. A synergy between site-specific and transient interactions drives the phase separation of a disordered, low-complexity domain. Proc. Natl. Acad. Sci. U. S. A. 120:e2305625120.

30. Oksuz, O., J.E. Henninger, R. Warneford-Thomson, M.M. Zheng, H. Erb, A. Vancura, K.J. Overholt, S.W. Hawken, S.F. Banani, R. Lauman, L.N. Reich, A.L. Robertson, N.M. Hannett, T.I. Lee, L.I. Zon, R. Bonasio, and R.A. Young. 2023. Transcription factors interact with RNA to regulate genes. Mol. Cell. 83:2449–2463.e13.

31. Kim, T.H., B.J. Payliss, M.L. Nosella, I.T.W. Lee, Y. Toyama, J.D. Forman-Kay, and L.E. Kay. 2021. Interaction hot spots for phase separation revealed by NMR studies of a CAPRIN1 condensed phase. Proc. Natl. Acad. Sci. 118.

32. Wang, J., J.-M. Choi, A.S. Holehouse, H.O. Lee, X. Zhang, M. Jahnel, S. Maharana, R. Lemaitre, A. Pozniakovsky, D. Drechsel, I. Poser, R.V. Pappu, S. Alberti, and A.A. Hyman. 2018. A Molecular Grammar Governing the Driving Forces for Phase Separation of Prion-like RNA Binding Proteins. Cell. 174:688–699.e16.

33. Fargason, T., N.I.U. De Silva, E. King, Z. Zhang, T. Paul, J. Shariq, S. Zaharias, and J. Zhang. 2023. Peptides that Mimic RS repeats modulate phase separation of SRSF1, revealing a reliance on combined stacking and electrostatic interactions. eLife. 12:e84412.

34. Kato, M., and S.L. McKnight. 2021. The low-complexity domain of the FUS RNA binding protein self-assembles via the mutually exclusive use of two distinct cross-β cores. Proc. Natl. Acad. Sci. 118:e2114412118.

