## Supplemental Figures for "Structure and position-specific interactions of prion-like domains in transcription factor Efg1 phase separation"

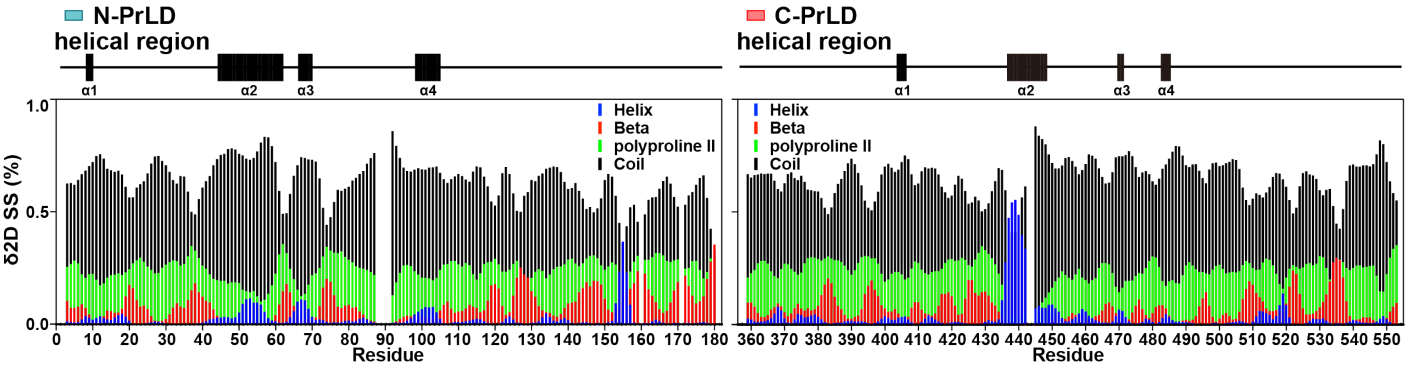


**Supplementary figure S1. Secondary structure analysis.** NMR chemical shifts based secondary structure prediction using the program δ2D indicates the formation of transient α-helices in both the N-PrLD and C-PrLD.

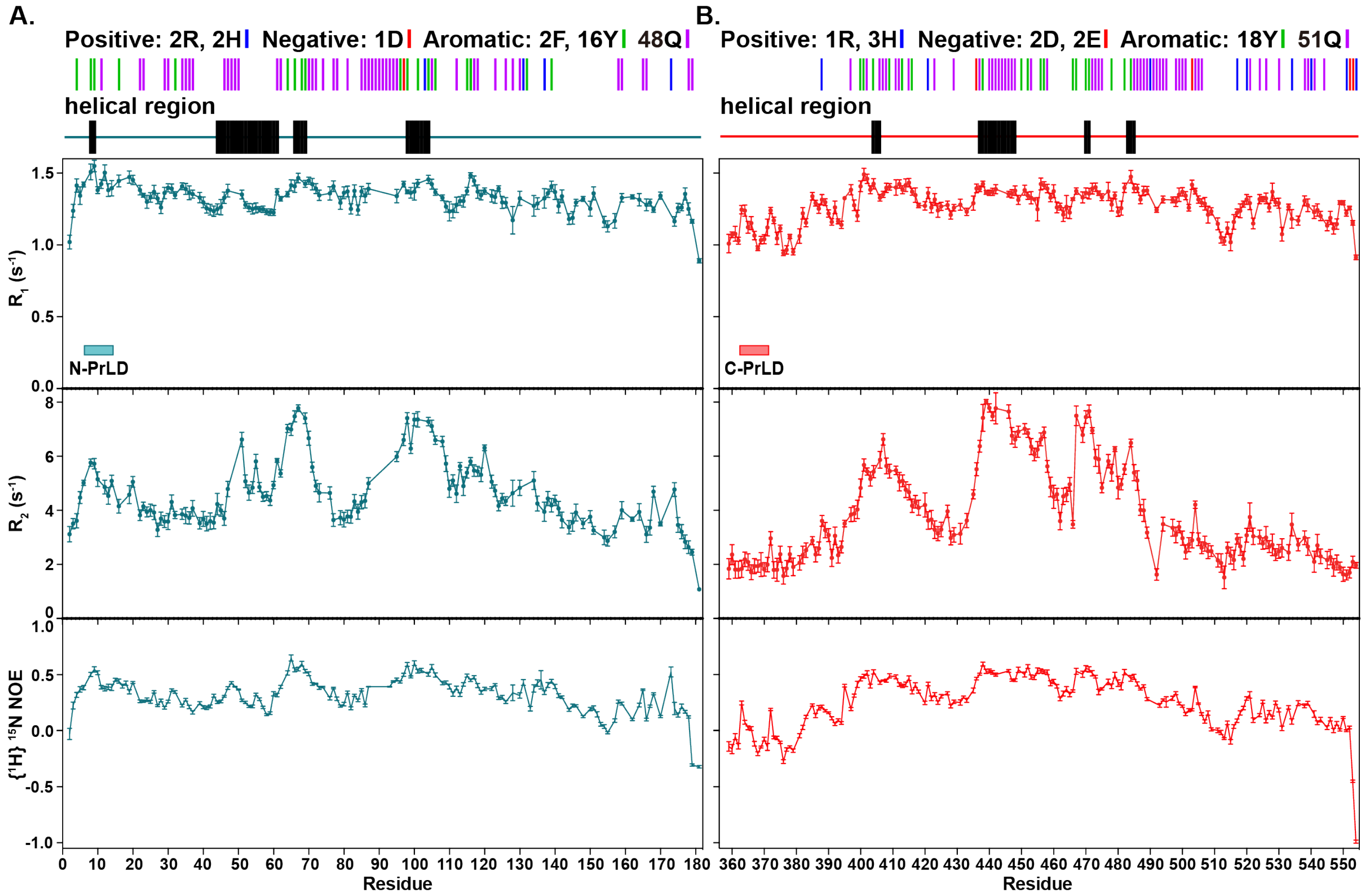


**Supplementary figure S2.** $^{15}$**N** $R_1$**,** $R_2$ **relaxation rate constant values and heteronuclear NOE measurements** for A) Efg1 N-PrLD and B) Efg1 C-PrLD in dilute solution (i.e. dispersed phase).

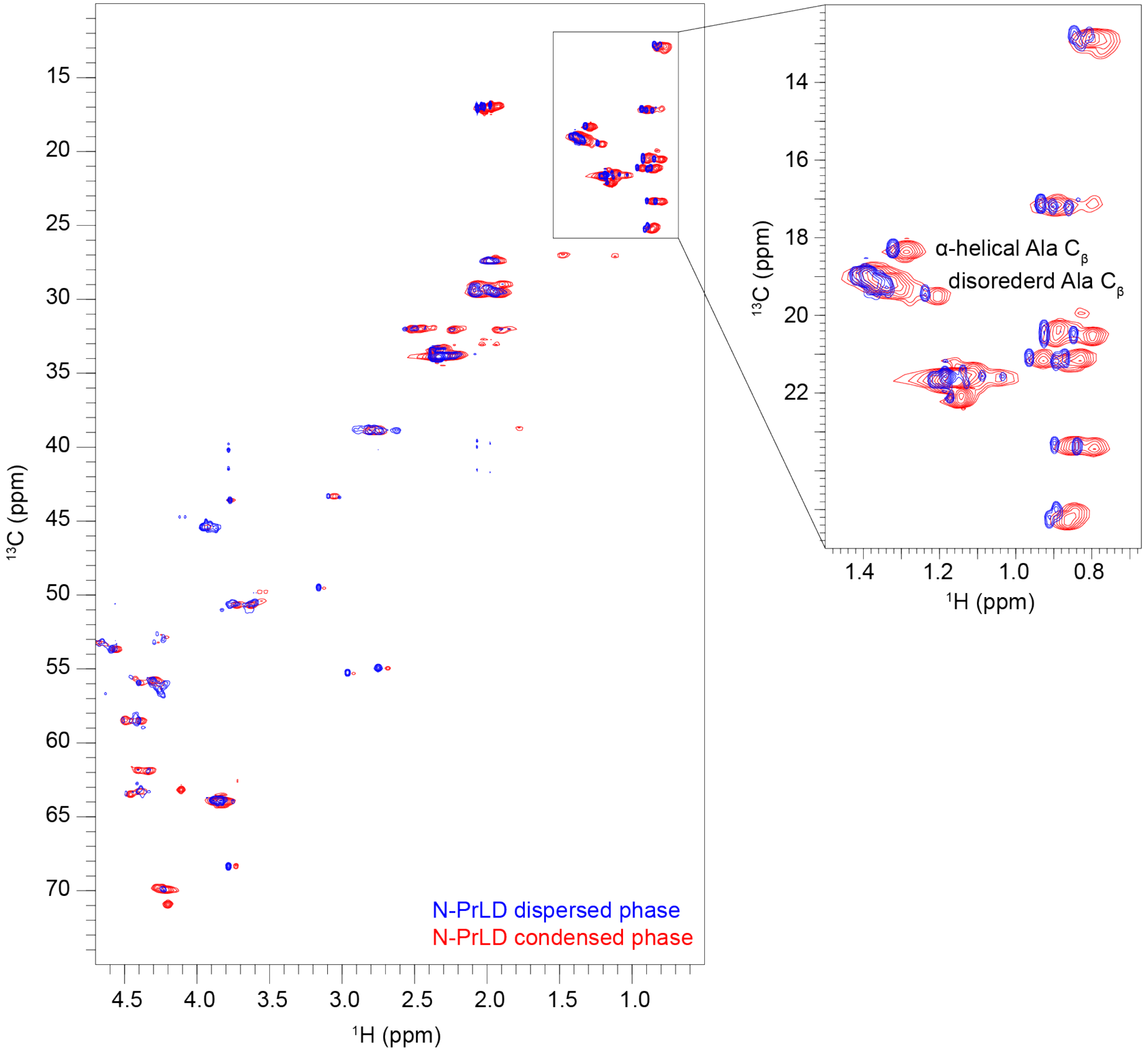


**Supplementary figure S3. $^{1}H$-$^{13}C$ HSQC spectra overlay of the dispersed phase and condensed phase Efg1 N-PrLD.** The well-overlapped peaks, especially on the $^{13}C$ dimension, suggest similar overall conformation in the two phases, including the α-helical alanine resonances found in both the dispersed and condensed phases.

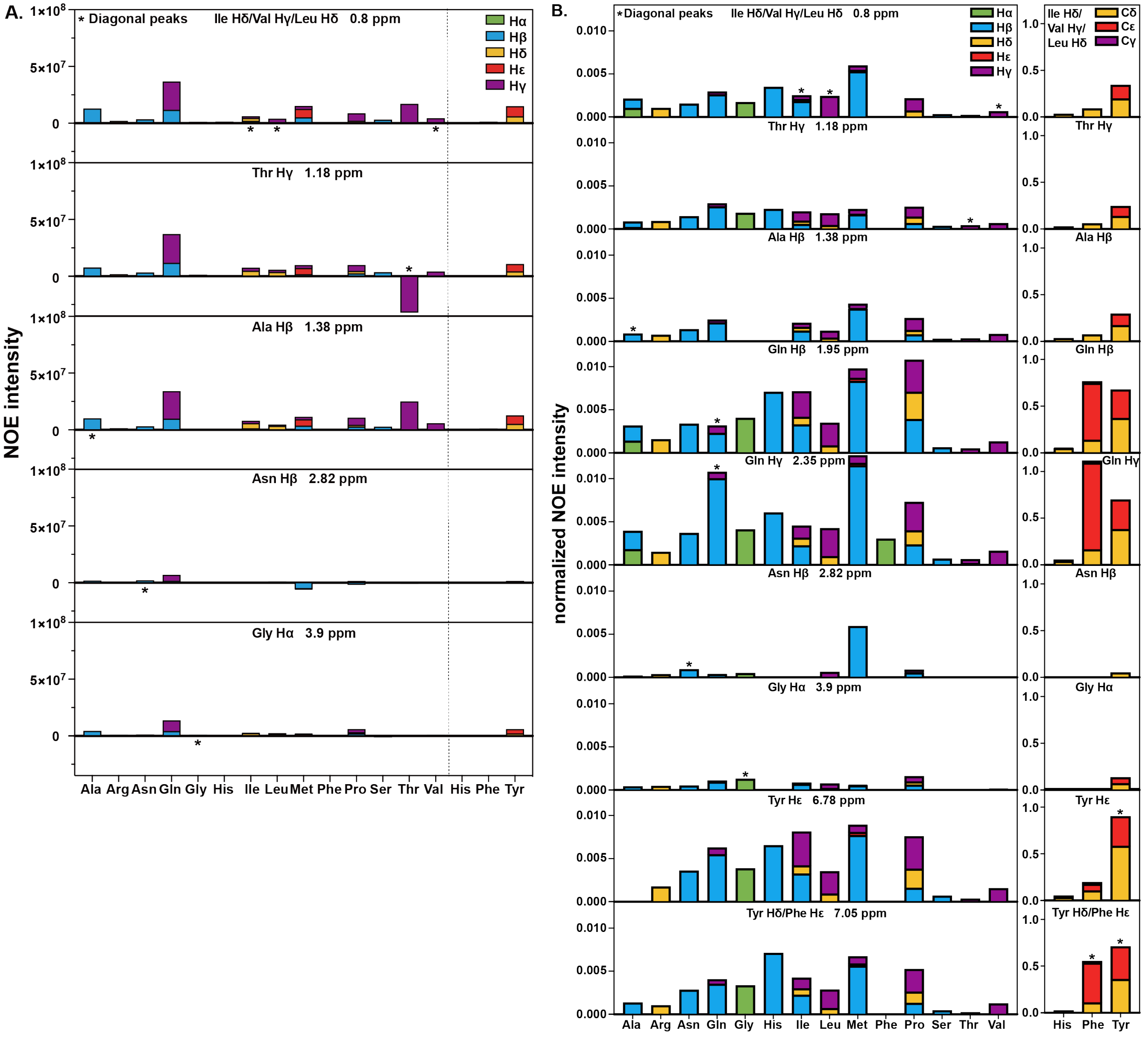


**Supplementary figure S4. Additional details for intermolecular NOE intensities.**

A) Quantification of additional spectra regions for NOE intensities from filtered/edited NOESY experiments.

B) NOE intensities normalized to the intensities of corresponding peaks in $^{1}H$-$^{13}C$ HSQC spectrum.

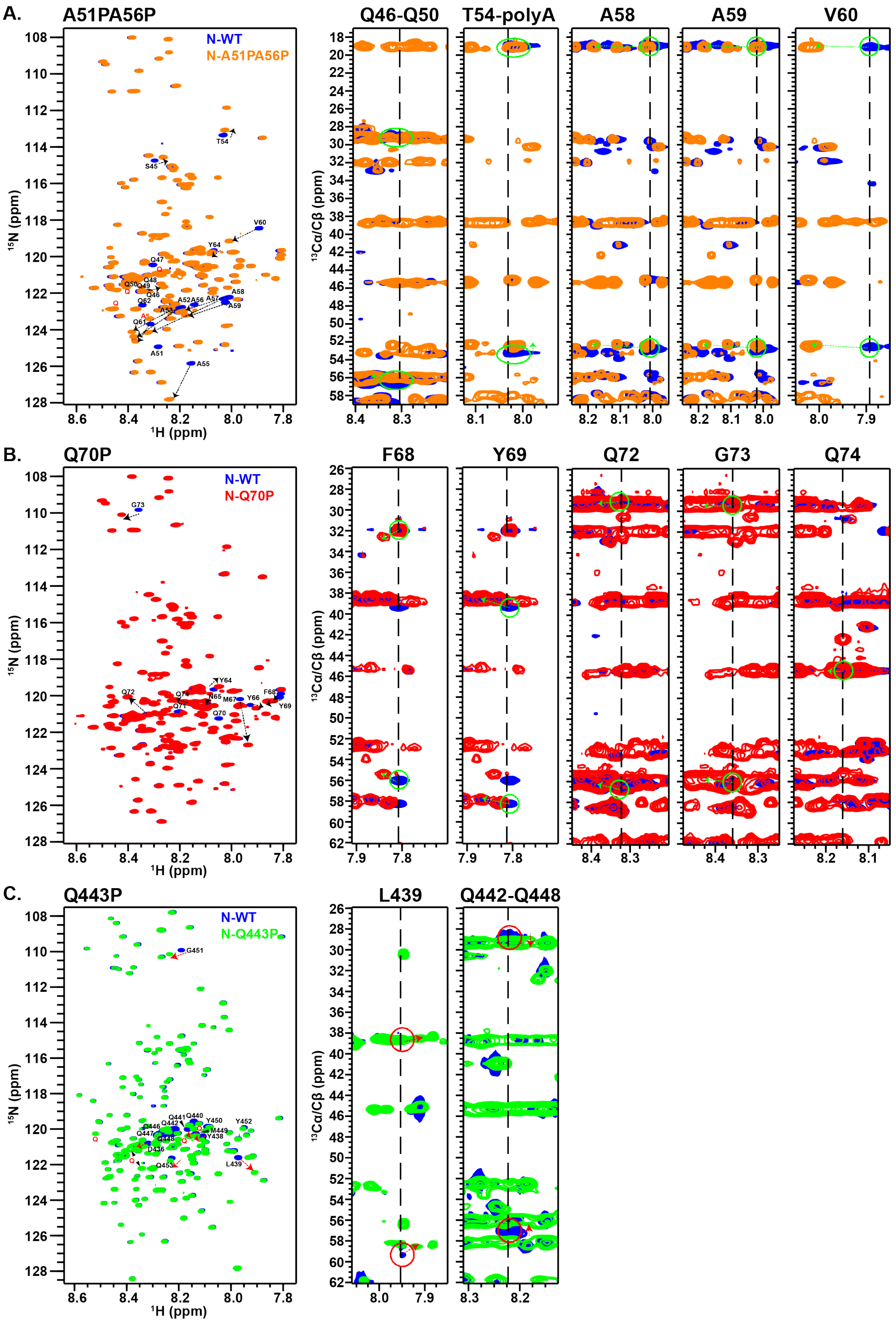


**Supplementary figure S5. Proline mutations disrupt helical structures in Efg1 N-PrLD and C-PrLD.** (A, B, C, left panels) $^{1}$H-$^{15}$N HSQC spectra show that when proline mutations were introduced, significant chemical shift perturbations were observed among residues within the same helix. The disruption of helical structures was also supported by $C_{\alpha}/C_{\beta}$ chemical shifts for residues nearby (A, B, C, right panels).
